# Alternative miRNAs: Human sequences misidentified as plant miRNAs in plant studies and in human plasma

**DOI:** 10.1101/120634

**Authors:** Kenneth W. Witwer

## Abstract

A recent study reported that “Plant miRNAs found in human circulating system provide evidences of cross kingdom RNAi” ^1^. Analysis of two human blood plasma sequencing datasets was said to provide evidence for uptake of plant miRNAs into human plasma. The results were also purportedly inconsistent with contamination ^1^. However, a review of these data suggests that they do not support dietary xenomiR uptake, but instead confirm previous findings that detection of rare plant miRNAs in mammalian sequencing datasets is artifactual. Only one putative plant miRNA (“peu-MIR2910) in this study mapped consistently above background, and this sequence is found in a human rRNA. Several other rarer but consistently mapped plant miRNAs also have 100% or near 100% matches to human transcripts or genomic sequences, and some do not map to plant genomes at all. These misidentified “alternative miRNAs”—including MIR2910 and MIR2911—emphasize the need for rigorous filtering strategies when assessing possible xenomiRNAs.

## INTRODUCTION

Reports of plant or other dietary miRNAs, or xenomiRs, entering mammalian circulation through the diet ^2–5^ generated excitement for the xenomiR transfer hypothesis, yet negative results of replication and reproduction studies have cast doubt on xenomiR transfer as a general mechanism ^6–12^. A prominent claim of xenomiR function ^2^ has also failed rigorous reproduction ^8^, unmasked as the result of an uncontrolled variable in the original experiment. Analyses of public datasets have revealed that studies of xenomiRs and other foreign-origin nucleic acids are fraught with artifacts: combinations of contamination, amplification or sequencing errors, permissive analysis pathways, and batch effects ^9^,^11^,^13–17^. A particularly comprehensive study recently found that foreign miRNAs in human biofluids and tissues do not match human food consumption, are marked by batch effects, and are thus most parsimoniously explained as artifacts ^14^. Studies of organisms with no exposure to plants have also found evidence of the same types of apparent plant contamination that plague some measurements of human samples ^9^,^18^. Liu et al ^1^ mapped sequencing data from two studies of human plasma and other samples ^19^,^20^ to various plant genomes using a 2010 plant miRNA database, PMRD ^21^, concluding that previous reports of dietary xenomiR transfer are supported. In this brief report, these results are examined critically.

## RESULTS

### Data evaluation

A cross-check of the source files and articles shows that the plasma data evaluated by Liu et al were from 198 plasma samples, not 410 as reported. Ninomiya et al sequenced six human plasma samples, six PBMC samples, and 11 cultured cell lines ^19^. Yuan et al sequenced 192 human plasma libraries (prepared from polymer-precipitated plasma particles) ^20^. Each library was sequenced once, and then a second time to increase total reads. Counts were presented as reads per million mapped reads (rpm) ^20^. In contrast, Liu et al appear to have reported total mapped reads in their data table ^1^. Yuan et al also set an expression cutoff of 32 rpm (log2 rpm of 5 or above). With an average 12.5 million reads per sample (the sum of the two runs per library), and, on average, about half of the sequences mapped, the 32 rpm cutoff would translate to around 200 total reads in the average sample as mapped by Liu et al ^1^.

### Only one putative plant miRNA above background levels

Consulting the Liu et al mapping table ^1^ and the Sequence Read Archive (SRA), results from duplicate sequencing runs from the Yuan et al dataset were combined, and two samples without reliable replicates were eliminated. A total of 1294 putative plant miRNAs had at least one mapped read in at least one of the remaining 190 samples. However, many of these miRNAs were identical orthologs or paralogs, and most were mapped at one or fewer rpm on average, and in only a small minority of samples. Across all samples, only one putative plant miRNA mapped above a median 200 read cutoff, roughly corresponding to the 32 rpm cutoff of Yuan et al (Table 1). All other RNAs, including previously reported xenomiRs such as MIR159a, MIR168a, and the plant ribosomal degradation fragment MIR2911 ^22–24^, were thus below the level of background noise established by the original investigators. Indeed, previously reported xenomiRs were mapped in few samples and below 1 rpm. The absence of these RNAs is confirmed by Liu et al’s analysis of the Ninomiya, et al study ^19^, where MIR159a, MIR168a, and MIR2911 mapped in none of the plasma samples. The single putative plant miRNA that mapped above background levels in this study was, again, peu-MIR2910 (Table 2).

**Table 1.** Summary of the most frequently mapping putative plant miRNAs in the Liu et al analysis of datasets from Yuan et al. Here, data from only 190 of 192 plasma samples were included, since all but the excluded 2 samples were successfully sequenced twice. miRNA inclusion criteria were: 1) Three total mapped reads according to Liu et al’s data in at least 10% of the samples and 2) discoverable putative mature sequence through miRBase, miRMaid, or miRNEST 2.0. An “estimated median rpm” value was calculated based on median total counts, average reads, and the midoint of the reported mapping percentage range. miRNAs with perfect human matches are in red, although most (see Table 1). Note that only MIR2910 consistently exceeds the rpm threshold set by Yuan et al.

| Putative miRNA | Samples w/<br>reads ≥3 | Average<br>total<br>counts | Median<br>total<br>counts | Max | Est. median<br>rpm in avg<br>sample |
| --- | --- | --- | --- | --- | --- |
| peu-MIR2910 | 190 | 1143.4 | 1072.5 | 2020 | 180.6 |
| peu-MIR2916 | 190 | 119.7 | 115 | 315 | 19.4 |
| tae-MIR2005 | 178 | 8.7 | 9 | 23 | 1.5 |
| peu-MIR2914 | 169 | 14.3 | 9 | 348 | 1.5 |
| tae-MIR2018 | 167 | 5.9 | 6 | 14 | 1.0 |
| ath-MIRf10482-akr | 161 | 7.3 | 6 | 29 | 1.0 |
| ppt-MIR896 | 147 | 4.6 | 4 | 15 | 0.7 |
| ptc-MIRf12412-akr | 47 | 2.7 | 2 | 18 | 0.3 |
| peu-MIR2911 | 42 | 2.6 | 2 | 10 | 0.3 |
| ppt-MIR894 | 39 | 2.3 | 2 | 18 | 0.3 |
| ptc-MIRf12524-akr | 28 | 1.8 | 1 | 5 | 0.2 |

**Table 2.** Putative plant miRNA mapping from the Ninomiya et al dataset. This dataset consisted of both cellular and plasma miRNA. Here, all results are shown in the left half of the table, and plasma results on the right. miRNA inclusion criteria were: 1) Three total mapped reads according to Liu et al’s data in at least 10% of the samples (cells and plasma together) and 2) discoverable putative mature sequence through miRBase, miRMaid, or miRNEST 2.0. “Avg rpm” is calculated from the total mapped reads and total reads per sample (not mapped reads). Putative miRNAs that met inclusion criteria in the Yuan et al study are italicized, and sequences with perfect human matches are in red.

| Cells (n=17) and plasma (n=6) |  |  |  |  |  | Plasma only (n=6) |  |  |  |  |
| --- | --- | --- | --- | --- | --- | --- | --- | --- | --- | --- |
| Putative miRNA | Samples<br>w/<br>reads<br>≥3 | Average | Median | Max | avg<br>rpm | Samples<br>w/<br>reads<br>≥3 | Average | Median | Max | avg<br>rpm |
| *peu-MIR2910* | 21 | 370 | 41.5 | 5369 | 21.3 | 6 | 1210 | 480.5 | 5369 | 83.2 |
| *ptc-MIRf10804-akr* | 17 | 26 | 26 | 60 | 1.2 | 0 |  |  | 0 | 0 |
| *ptc-MIRf12412-akr* | 17 | 63 | 61 | 156 | 2.8 | 0 |  |  | 0 | 0 |
| *tae-MIR2018* | 17 | 39 | 29 | 163 | 1.7 | 0 |  |  | 0 | 0 |
| *ptc-MIRf12524-akr* | 10 | 13 | 10.5 | 31 | 0.4 | 2 | 16 | 16 | 31 | 0.4 |
| *tae-MIR2005* | 7 | 17 | 11 | 69 | 0.4 | 0 |  |  | 0 | 0 |
| *ath-MIRf10045-akr* | 4 | 5 | 4 | 10 | 0.1 | 0 |  |  | 0 | 0 |
| *ath-MIRf10046-akr* | 4 | 5 | 4 | 10 | 0.1 | 0 |  |  | 0 | 0 |
| *peu-MIR2914* | 3 | 26 | 11 | 81 | 0.3 | 3 | 33.7 | 19 | 81 | 1.2 |
| *peu-MIR2915* | 3 | 5 | 4.5 | 8 | 0.0 | 0 |  |  | 0 | 0 |

### Lowering the threshold: still only a handful of possible xenomiRs

Since only one plant miRNA appeared to map consistently above background, the inclusion threshold of Yuan et al was relaxed to include all miRNAs with three or more mapped reads (Liu et al data) in 10% or more of the samples from either study. These are rather permissive criteria but may at least screen out some false positives due to amplification and sequencing errors. All samples from the Ninomiya study were included, despite the fact that most were not plasma. 11 miRNAs satisfied these criteria for the Yuan et al data (Table 1). (One low-mapping miRNA was excluded because its sequence could not be found in miRBase ^25^,^26^, miRMaid ^27^, miRNEST 2.0 ^28^ or indeed through any searches attempted.) 10 satisfied the criteria from the Ninomiya study (Table 2), including one sequence that was part of another (compare ath-MIRf10046-akr and ath-MIRf10045-ak, Table 3). However, if only the plasma samples from the latter study are considered, three miRNAs remain (Table 1). In total, 15 putative miRNAs satisfied the permissive inclusion criteria, including five (Yuan only), four (Ninomiya only), and six (both) (Table 3).

**Table 3.**
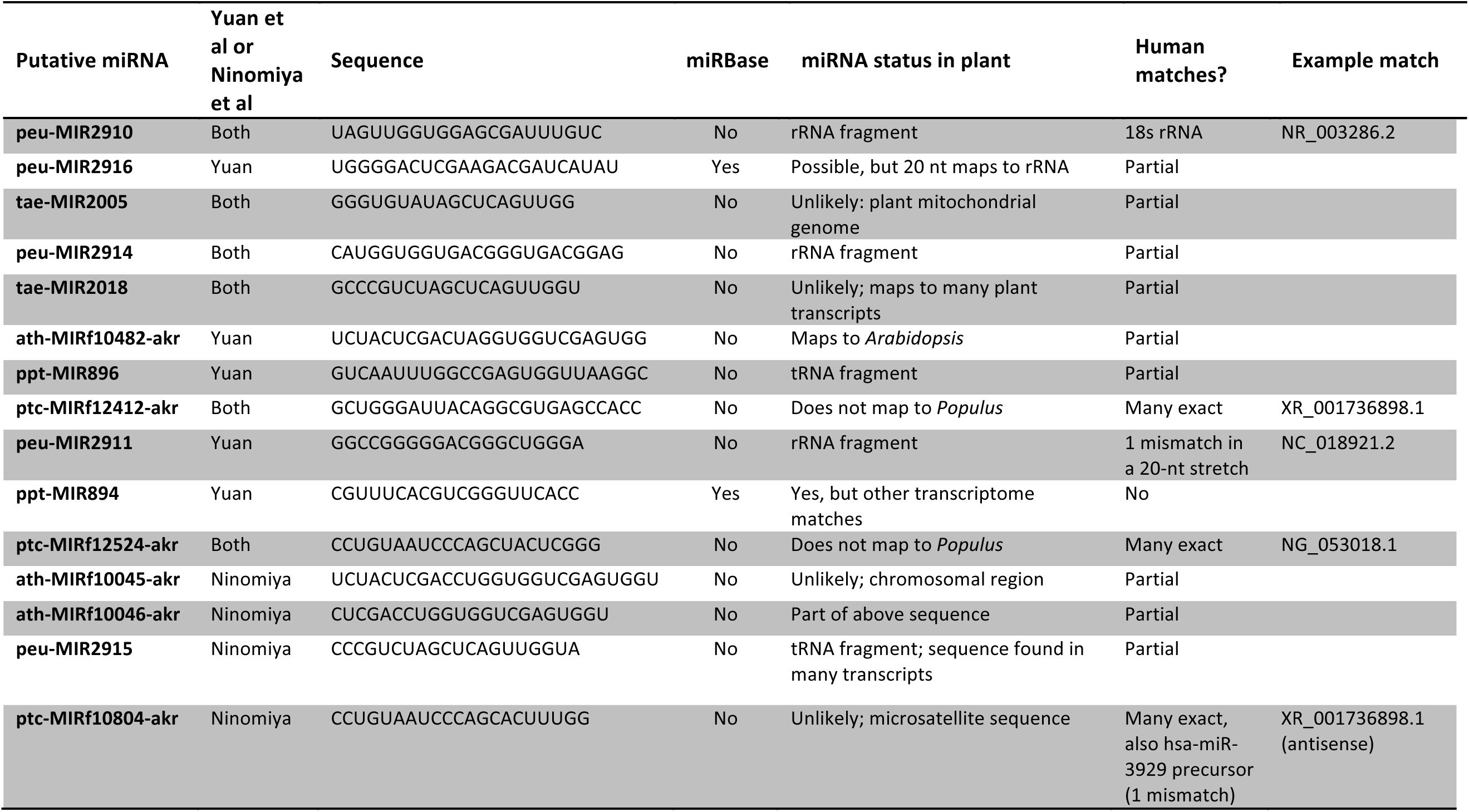
Putative plant miRNAs mapped by Liu et al from the Yuan et al or Ninomiya et al studies. Inclusion criteria were: 1) Three total mapped reads according to Liu et al’s data in at least 10% of the samples in the respective studies and 2) discoverable putative mature sequence through miRBase, miRMaid, or miRNEST 2.0. miRNA status was considered unlikely if miRBase listed the miRNA as a non-miRNA or if the sequence mapped to non-miRNA regions. Human matches were exact (with an example given), “partial” (at least 15 nt stretches with 100% identity), or as otherwise described. Note that the ath-MIRf10046-akr is found within the ath-MIRf10045-akr sequence.

### To miR or not to miR

As miRNA discovery, validation, and annotation has advanced, numerous reported miRNAs have been reclassified as degradation fragments of other noncoding RNAs (ncRNAs). A classic example is MIR2911, a plant rRNA degradation fragment that has been misidentified as a microRNA. Interestingly, only 2 of the 15 miRNAs identified as plant miRNAs in this study are annotated in miRBase. Although some of these sequences may represent rare or unusually structured miRNAs, several are part of non-miRNA ncRNAs or other sequences that seem unlikely, at least at first glance, to give rise to microRNAs. Among the apparently misidentified miRNAs is MIR2910, the most abundant plant miRNA identified by Liu et al. The MIR2910 sequence, UAGUUGGUGGAGCGAUUUGUC, is found in the highly conserved and expressed large subunit (LSU) rRNA of plants, and has been specifically removed from miRBase as a non-miRNA. Even the two identified miRNAs that remain in miRBase, MIR2916 and MIR894, are not above question. A 20 nucleotide stretch of MIR2916 map to rRNA, while the full MIR894 sequence appears to be found in a variety of plant transcripts.

### Human sequences in the plant database and vice-versa

Curiously, several sequences did not map to the species to which they were ascribed by the PMRD ^21^. Unfortunately, the PMRD could not be accessed directly during this study; however, other databases appear to provide access to its contents. Specifically, ptc-MIRf12412-akr and ptc-MIRf12524-akr did not map to *Populus* or to other plants. The poplar tree is also not a common dietary staple of human populations. In contrast, both sequences mapped with 100% identity and coverage to numerous human sequences (Table 3). ptc-MIRf10804-akr had numerous 100% identity human matches, plus a 1-mismatch alignment to the human miR-3929 precursor. Other miRNAs, including MIR2911, also displayed some lesser degree of matching to human transcripts or the genome. Strikingly, the putative MIR2910 sequence is not only a fragment of plant rRNA; it has a 100% coverage, 100% identity match in the human 18S rRNA (see NR_003286.2 in GenBank; Table 3). These matches of putative plant RNAs with human sequences are difficult to reconcile with the statement of Liu et al that BLAST of putative plant miRNAs “resulted in zero alignment hit” ^1^, suggesting that perhaps a mistake was made, and that the BLAST procedure was performed incorrectly.

## CONCLUSION

In mammalian studies, mapping of MIR2910 and other dubious plant miRNAs is best explained as mapping of human degradome fragments to plant RNAs that are in some cases genuine sequences but not miRNAs, and in other cases, human sequences that have contaminated plant RNA samples and databases. Re-analysis of the results of Liu et al ^1^ thus echoes the recent findings of Kang, Bang-Berthelsen, and colleagues ^14^, as well as previous negative findings surrounding dietary xenomiRs, summarized above. A stringent data analysis procedure, such as filtering all reads against the ingesting organism genome/transcriptome with one or two mismatches, then requiring perfect matches of remaining reads against plant or other foreign organisms, would engender higher confidence that “foreign” RNAs are not simply amplification or sequencing artifacts. Indeed, pre-mapping to the ingesting organism’s genome may not be sufficient; as shown ^14^, the largest number of xenomiRs in some human studies are from rodents, likely because of proximity in research laboratories. Therefore, it may be best to screen against mammalian sequences in general, and perhaps also against widespread microbe contaminants. Of course, even the most stringent analysis procedures cannot distinguish a physical contaminant from a “real” read; therefore strict process controls are also needed to assess possible contamination. In general, such controls have not been done in existing studies.

This report underlines the danger in assuming that xenomiRs in mammalian material originate from the diet. When the species and roles are reversed—for example, with the finding of human sequences in a list of poplar tree miRNAs—few analysts would conclude that poplar trees consume humans. The simplest explanation is that the sequenced plant material was contaminated with human nucleic acid. In the same way, the extremely low-level, variable, and batch-effect prone concentrations of several plant sequences in human plasma and tissue could be due to uptake from the diet, albeit at levels far too low to affect physiologic processes. However, artifact remains the simplest explanation.

## METHODS

Plant mapping results from Liu et al ^1^ (total mapped counts) were downloaded from the BMC Genomics website. Accession numbers of sequencing datasets were checked against the publications of Ninomiya et al ^19^ and Yuan et al ^20^, as well as the Sequence Read Archive (SRA, https://www.ncbi.nlm.nih.gov/sra). Data were sorted and analyzed in Microsoft Excel. Plant miRNA sequences were obtained from miRBase (http://mirbase.org/) ^29^. Because certain plant sequences have been removed from miRBase because they have been identified as ncRNA degradation artifacts, the plant microRNA database (PMRD) ^21^ was consulted; however, repeated attempts to access the site (http://bioinformatics.cau.edu.cn/PMRD) were unsuccessful, so information was retrieved instead from miRMaid (http://140.mirmaid.org/home) ^27^ or miRNEST 2.0 (http://rhesus.amu.edu.pl/mirnest/copy/home.php) ^28^. All analysis files are available on request.

